# Divisive normalization in mouse V1 is unchanged by increased thalamocortical short-term depression

**DOI:** 10.64898/2026.07.13.738133

**Authors:** J. Leonie Cazemier, Louis R.W.M. Douin, L. Niels Cornelisse, Mohit Dubey, Maarten H.P. Kole, J. Alexander Heimel

## Abstract

Divisive normalization describes how neuronal responses are scaled by the activity of a broader pool of inputs and accounts for several nonlinear response properties in primary visual cortex (V1). It accurately describes contrast gain control, as well as the response to overlapping gratings. Nevertheless, how this computation is performed by the brain is unclear. One mechanism suggested to underlie divisive normalization in V1 is short-term depression (STD) at thalamocortical synapses. Here, we used a Vglut2-Cre dependent heterozygous deletion of Stxbp1 in mice as an *in vivo* model for increased STD. This enabled us to test whether STD is responsible for divisive normalization in V1. As expected, responses were shorter in the mutant mice compared to wild-type siblings. Although we did find other differences in visual responses, notably in direction selectivity and response linearity, neither the contrast response function, nor the amount of cross orientation suppression differed between the two groups. We conclude that STD is likely not involved in divisive normalization of visual responses in mouse V1. Instead, short-term plasticity shapes the temporal integration through which selective cortical responses are constructed.

## Introduction

Neuronal responses to sensory stimuli are strongly influenced both by stimulus history and by the presence of other stimuli. Stimulus history can produce adaptation, shifting the response range to match recent input conditions (Barlow, 1961; Ferguson & Cardin, 2020; Schwartz & Simoncelli, 2001). A related but distinct computation is divisive normalization, in which the response to a stimulus is scaled by a normalization term reflecting pooled activity across a population of filters (Carandini & Heeger, 2012; Heeger, 1992). This computational model successfully describes several non-linear properties of sensory responses, such as contrast saturation, contrast adaptation, cross-orientation suppression (Heeger, 1992), dependence of visual acuity on contrast (Heimel et al., 2010), attention (Reynolds & Heeger, 2009) and multisensory integration (Ohshiro et al., 2017), in rodents, carnivores, non-human primates, and humans. However, its cellular mechanism is not well understood (Carandini & Heeger, 2012; Heeger, 1992).

One mechanism suggested to underlie divisive normalization is short-term synaptic depression (STD). STD is a reduction in the post-synaptic response during high-frequency pre-synaptic firing (Wu & Borst, 1999; Zucker & Regehr, 2002). It takes place at the range of tens of milliseconds to seconds (Campagnola et al., 2022; Zucker & Regehr, 2002) and creates a non-linear gain function (Abbott & Regehr, 2004; Deng & Klyachko, 2011; Rothman et al., 2009). STD can act as gain control (Abbott et al., 1997; Chung et al., 2002; Tsodyks & Markram, 1997), and STD in thalamocortical synapses can produce the saturation of responses in the primary visual cortex (V1) with increasing contrast that is described by divisive normalization (Chance et al., 1998; Kayser et al., 2001) (**Fig. 1A**). Furthermore, STD of these thalamo-cortical synapses has been proposed to underlie cross-orientation suppression in V1 (Carandini et al., 2002; Freeman et al., 2002). Cross-orientation suppression occurs when the response to two overlaid orthogonal gratings is lower than the linear summation of responses to the individual gratings. The origin of this phenomenon was long thought to be intracortical GABAergic inhibition (Morrone et al., 1987), but later work showed this to be incorrect because even masking gratings that do not elicit a response in V1 because of a high drift speed or adaptation, can suppress responses to a test grating (Barbera et al., 2022; Freeman et al., 2002). Earlier models showed that contrast-dependent nonlinearities and responses to multiple gratings could arise from local V1 circuitry and synaptic depression (Kayser et al., 2001; Lauritzen et al., 2001). Carandini and colleagues then proposed that STD in thalamocortical synapses could elicit cross-orientation suppression itself (Carandini et al., 2002; Freeman et al., 2002; Mrsic-Flogel & Hübener, 2002) (**Fig. 1B**).

**Figure 1.**
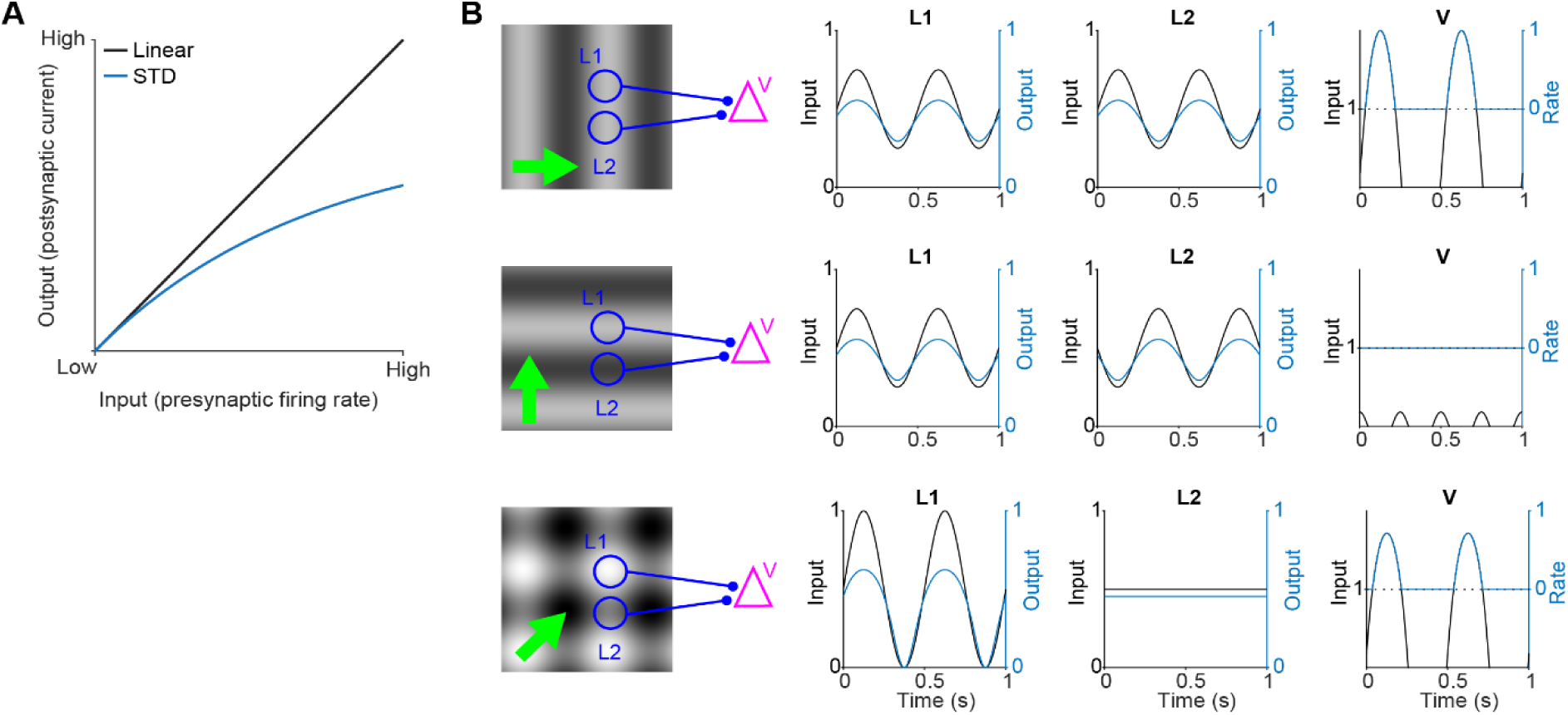
Models explaining the putative role of STD in divisive normalization. **(A)** STD could be responsible for non-linear gain in contrast response functions. With STD, the transmission efficiency decreases as the presynaptic firing rate increases. **(B)** Model of how STD could be responsible for cross-orientation suppression in V1. Left: Visual stimulus with overlaid two blue circles representing the receptive fields of two Lateral Geniculate Nucleus (LGN) neurons (L1/L2). Green arrows represent direction of motion of the stimulus across the receptive fields. V represents response of a V1 neuron. Top row: during the optimal stimulus, LGN neurons L1 and L2 fire in phase. The summed normalized input crosses the firing threshold in V. Middle row: during the orthogonal stimulus, the LGN neurons fire out of phase, and the firing threshold in V is not reached. Bottom row: L1 fires strongly as it is exposed to the varying luminance of the drifting plaid. L2 fires at a steady rate as it is not exposed to any luminance change. The summed normalized input in V is smaller for the plaid stimulus than for the optimal stimulus, because of the extra synaptic depression of the L1-to-V synapse from the orthogonal grating.

Thalamocortical synapses can already be tonically depressed by ongoing activity in vivo, which may limit the additional depression evoked by visual stimulation (Boudreau & Ferster, 2005; Reig et al., 2006). Because these recordings were performed under anesthesia, the strength and impact of thalamocortical depression may differ in awake animals (Reinhold et al., 2015; Swadlow et al., 2002). Whether changing the strength of short-term depression alters contrast saturation or cross-orientation suppression in V1 has not been tested directly.

Here, we tested the role of short-term depression in divisive normalization using a Vglut2-Cre-dependent heterozygous deletion of Stxbp1 (**Fig. 2A**). Stxbp1, also known as Munc18-1, is a presynaptic protein involved in synaptic vesicle release and recovery (Han et al., 2010; Stepien et al., 2019; Verhage et al., 2000; Voets et al., 2001), and its heterozygous deletion increases short-term depression without substantially affecting baseline transmission (Miyamoto et al., 2019a; Toonen et al., 2006). We found that increased short-term depression in white-matter inputs to layer 4 shortened visual responses in V1, but left contrast saturation and cross-orientation suppression unchanged. In contrast, direction selectivity and response linearity were altered, indicating that altering short-term plasticity changes temporal integration and feature selectivity of V1 responses, but does not change divisive normalization.

**Figure 2.**
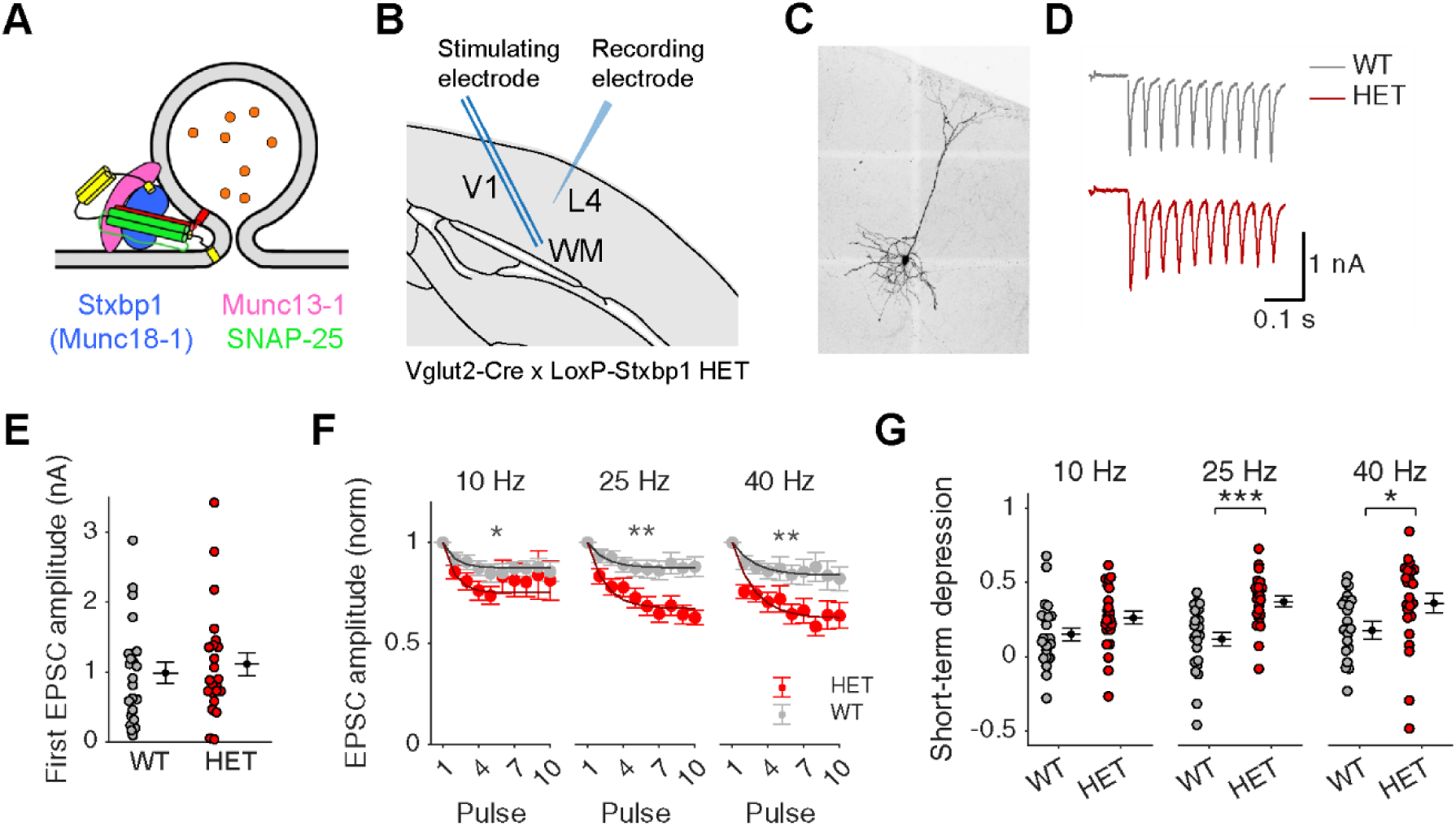
**Increased STD in V1 neurons of mice carrying a heterozygous deletion of Stxbp1 in Vglut2^+^ neurons**. **(A)** Schematic drawing indicating the function of Stxbp1 in synaptic vesicle docking and fusion (adapted from (Stepien et al., 2019). **(B)** A bipolar stimulating electrode is placed in the white matter (WM) of a coronal slice of V1 and a neuron in L4 is recorded using whole-cell patch voltage clamp. **(C)** Example of recorded and stained L4 neuron. **(D)** Example current traces to 10 stimulation pulses at 25 Hz. Stimulation artefact has been removed. **(E)** No difference in the amplitude of the EPSC in response to the first pulse in Vglut2-Stxbp1 WT and HET mice. Markers indicate neurons. Level and error bar indicate mean and s.e.m. **(F)**. More decrease of EPSC amplitude during pulse trains in Vglut2-Stxbp1 HET mice for 10, 25, and 40 Hz. Curves are non-linear mixed effect model fits of exponential decays with an variable offset and equal decay constants per stimulation frequency. *: p<0.05; **: p<0.01; ***: p<0.001. **(G)** Increased STD in Vglut2-Stxbp1 HET mice. Detailed statistics are in **Supp. Table 1**.

## Results

We used a Vglut2-Cre dependent heterozygous Stxbp1-deletion strategy to test whether short-term depression of the excitatory thalamocortical synapses plays a role in divisive normalization. Short-term synaptic depression is increased when Stxbp1 is heterozygously knocked out, as has been shown for Hippocampal neurons *in vitro* (Toonen et al., 2006) and cortical-striatal connections *in vivo* (Miyamoto et al., 2019a). First, we investigated whether short-term plasticity is indeed altered in our model. To achieve this, we made coronal slices of V1, placed a bipolar stimulation electrode in the white matter and performed whole-cell patch clamp recordings of excitatory neurons in layer 4, the layer receiving its main drive from the dLGN (**Fig. 2B-C**). There were no differences in series resistance, capacitance or resting membrane potential (**Supp. Fig 1**, **for details of all statistics of this study, see Supp. Table 1**). We gave repeated pulse trains of 10 pulses, at 10, 25 and 40 Hz (**Fig. 2D**). The first EPSC amplitude was not different in neurons from HET mice or WT mice (p = 0.54, **Fig. 2E**), confirming that baseline transmission is unaffected by the heterozygous knock-out of Stxp1, as expected from previous work (Miyamoto et al., 2019b; Toonen et al., 2006). The amplitude for EPSC in response to later stimuli in the train goes down to a lower level in HET mice than in WT mice at tested pulse train frequencies (10 Hz: p = 0.037; 25 Hz: p = 0.0013; 40 Hz: p = 0.0094, **Fig. 2F**). The amount of short-term depression, the relative reduction from the first to the last EPSC in the train, is higher in neurons of HET mice than in WT mice (10 Hz: p = 0.051; 25 Hz: p = 0.00017; 40 Hz: p = 0.017). The Stxbp1 deficieny thus results in an increased short-term depression.

Next, we performed in vivo extracellular recordings of neural responses from V1 of urethane-anesthetized mice (**Fig. 3**). We used a stimulus displaying small white squares in random positions on a black background (**Fig. 3A**). Using this stimulus we were able to investigate some basic properties of V1 of our knockout model. The distribution of receptive field sizes did not differ between Vglut2-Stxbp1 WT and HET mice (**Fig. 3C**; p = 0.95). This is an indication that the basic spatial input properties of V1 are unchanged. When we look into the responses of neurons to the luminance change in their preferred patch location, however, we see that the firing rates for HET mice are different compared to those of the WT mice (**Fig. 3D-E**). The peak rates were not different between WT and HET mice (**Fig. 3F,** p = 0.22), but the half-life of the responses was significantly shorter for HET mice than for WT mice (**Fig. 3G,** p = 0.005), in line with earlier patch recordings measuring short-term depression in synapses in which Stxbp1 was heterozygously knocked-out (Miyamoto et al., 2019a; Toonen et al., 2006). From this, we can conclude that short-term depression of visual responses was increased upon the heterozygous deletion of Stxbp1 in Vglut2-expressing synapses.

**Figure 3.**
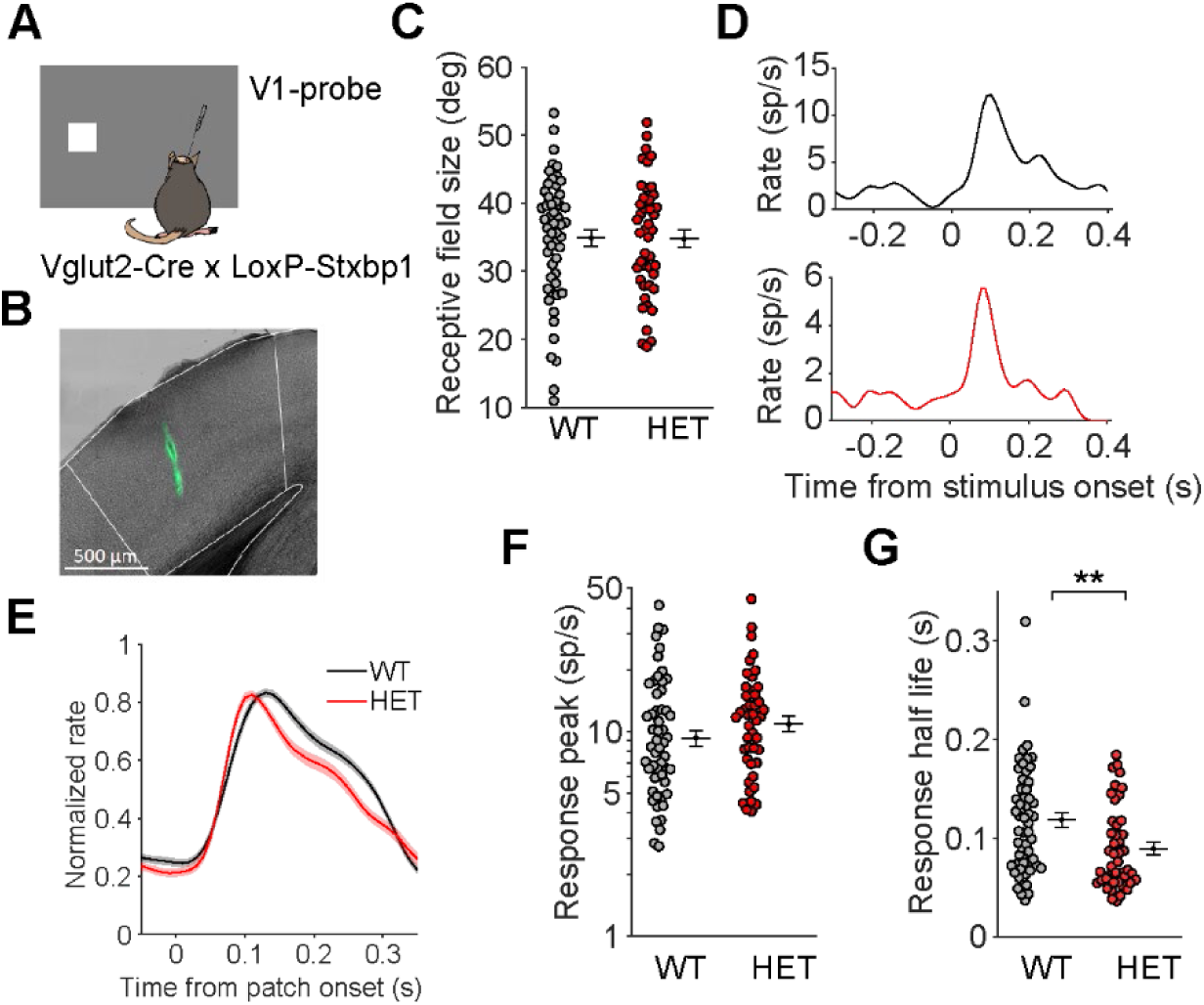
**V1 neurons in Vglut2-Stxbp1 HET mice show similar peak rates but shorter visual responses**. **(A)** *In vivo* recording in V1 of Vglut2-Stxbp1 HET mice. **(B)** Histological confirmation of the recording site in V1. Greyscale: brightfield. Green: Neuro-diO. **(C)** No difference in receptive field size between Vglut2-Stxbp1 WT and HET mice. Dots indicate units (7 WT mice, 5 HET mice). Error bars indicate mean ± s.e.m. **(D)** Examples of responses to white square in the receptive field center of a neuron of a WT (top) and HET mouse (bottom). **(E)** Population average PSTHs**. (F)** Peak response amplitudes are not different. **(G)** Response half-life is shorter in Vglut2-Stxbp1 HET mice. **: p<0.01. Detailed statistics are in **Supp. Table 1**.

Having established that V1 responses to static stimuli in our HET mice indeed have a shorter half-life than those of WT mice, we next turned to investigate divisive normalization in V1 through the contrast gain function. We first assessed the preferred direction of the recorded neurons to investigate the contrast tuning of V1 neurons at their preferred direction. Different aspects of the contrast tuning can be quantified by the curve’s dynamic range and the C50 (**Fig. 4A**). The dynamic range is the range of contrasts between which the neuron fires between 25% and 75% of its maximal firing rate. The C50 is the contrast at which the neuron fires at 50% of its maximal rate. The contrast tuning curves of Vglut2-Stxbp1 WT and HET mice look almost the same (**Fig. 4B-C**). Although contrast-response functions in mouse V1 are often less strongly saturating than those described in cat or primate V1 (King et al., 2016), the examples in **Fig. 4B** show that our sample included both neurons with clear response saturation and neurons with more nearly linear contrast-response profiles. Both the C50 (**Fig 4D**, p = 0.11) and the dynamic range (**Fig. 4E**, p = 0.83) are not significantly different between WT and HET mice. This indicates that the contrast gain function in V1 is not affected by the altered short-term plasticity in the HET mice.

**Figure 4.**
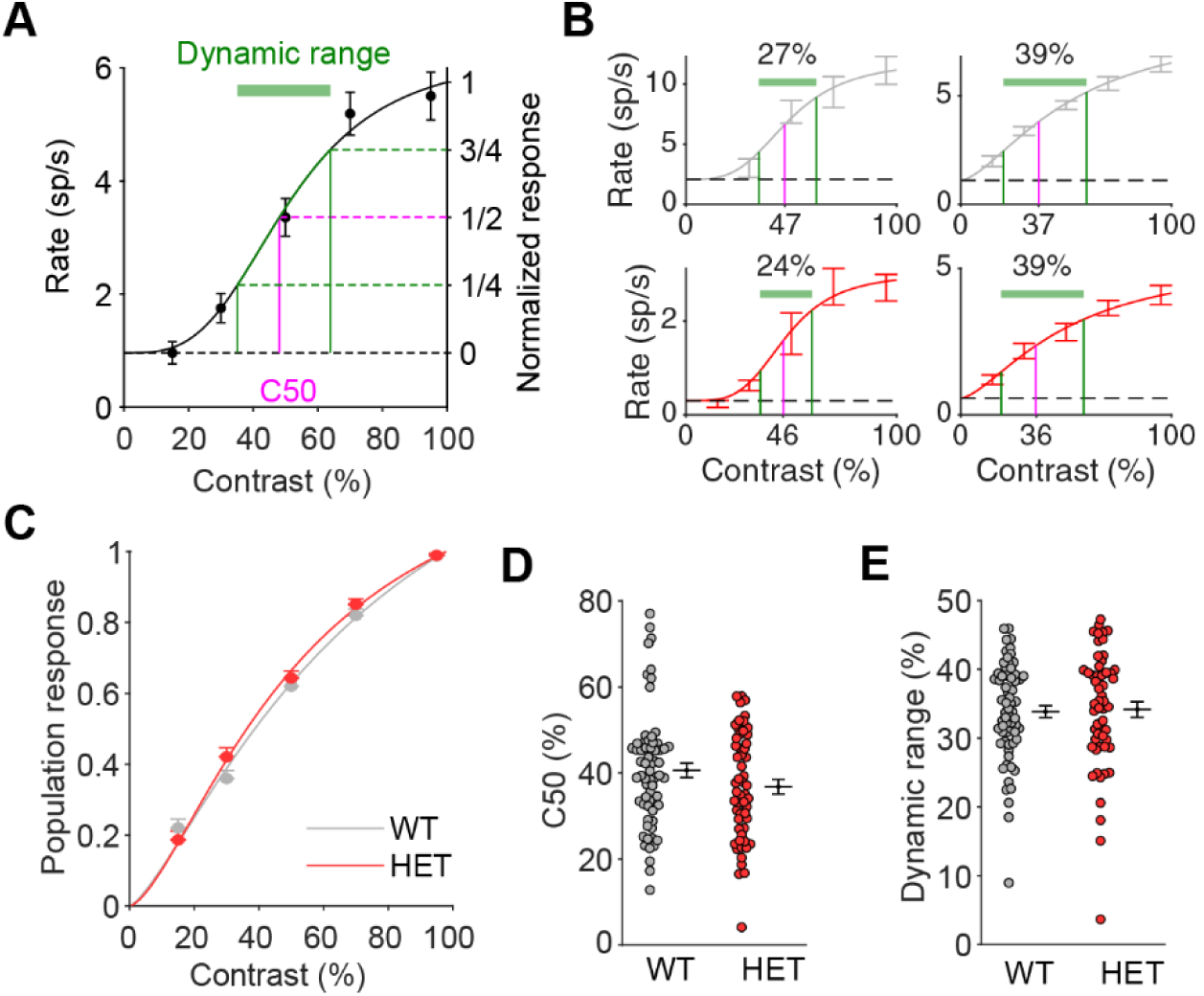
No difference in contrast tuning in Vglut2-Stxbp1 HET vs. WT mice. **(A)** Contrast tuning curve with the dynamic range indicated in green and C50 indicated in pink. **(B)** (B) Example contrast tuning of neurons in Vglut2-Stxbp1 WT (grey) and HET (red) mice, showing both neurons with clear response saturation and neurons with more nearly linear contrast-response profiles. **(C)** Population average contrast tuning curves. Lines show Naka-Rushton fits. Error bars indicate mean ± S.E.M.. **(D)** No difference in C50. Dots indicate neurons (7 WT mice, 5 HET mice). Error bars indicate mean ± s.e.m. **(E)** No difference in dynamic range. Detailed statistics are in **Supp. Table 1**.

As a second test for the role of STD in divisive normalization, we next tested whether cross-orientation suppression is affected in our knockout model. To do this, we used 3 different drifting grating stimuli (**Fig. 5A**): the optimal direction, the orthogonal direction (each at 50% contrast), and a 100% contrast plaid generated by computing the linear sum of the difference of each of the gratings compared to the mean luminance, added to the mean luminance. Since we only performed this experiment using the population-optimal stimuli, we picked the optimal stimulus for each neuron as the direction to which the neuron responded most in this test. To quantify the responses to these stimuli, we computed a cross-orientation suppression index (XOSI)

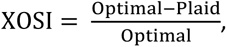

which indicates how much the response to the optimal grating is suppressed by superposition of the cross-orientation grating, as well as a masking index (MI; (Barbera et al., 2022; Guan et al., 2020)

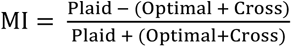

which indicates the linearity of the responses. An MI larger than zero means that the responses to the two gratings add superlinearly when they are superposed. **Figure 5B** shows the responses to each of the stimuli of two example neurons. The response of the left neuron to the plaid stimulus is smaller than the response to the optimal grating. This leads to a positive XOSI, as the neuron is suppressed by adding the cross-orientation, and a negative MI because the plaid response is also smaller than the summed responses to the optimal and cross orientations. The second example neuron shows a much higher response to the plaid than to the optimal stimulus, higher even than the sum of the optimal and cross responses – a superlinear addition. Hence, its XOSI is negative and its MI is positive. When we investigate the population responses to the cross and plaid stimulus, normalized to the optimal stimulus (**Fig. 5C-D**), we see a mix of responses: some neurons show cross-orientation suppression, whereas most neurons show some form of addition and some show cross-orientation facilitation. This mix of suppression and facilitation is consistent with previous findings in mouse V1 (Barbera et al., 2022; Muir et al., 2015; Palagina et al., 2017). The distribution of masking indices (−0.14 ± 0.02, mean ± s.e.m.) was similar to the spiking data recently reported in anesthetized mouse V1 (Barbera et al., 2022). The populations are similar between the Vglut2-Stxbp1 WT and HET mice: neither the XOSI (**Fig. 5E,** p = 0.73), nor the MI (**Fig. 5F,** p = 0.59) differs between the two groups of mice. We conclude that, similar to the contrast gain control, the extent of cross-orientation suppression in V1 is not affected by increased short-term depression. This finding fits with recent work suggesting that cross-orientation interactions in mouse V1 arise largely from feedforward thalamic input, leaving limited need for an additional nonlinear contribution at the thalamocortical synapse (Barbera et al., 2022).

**Figure 5.**
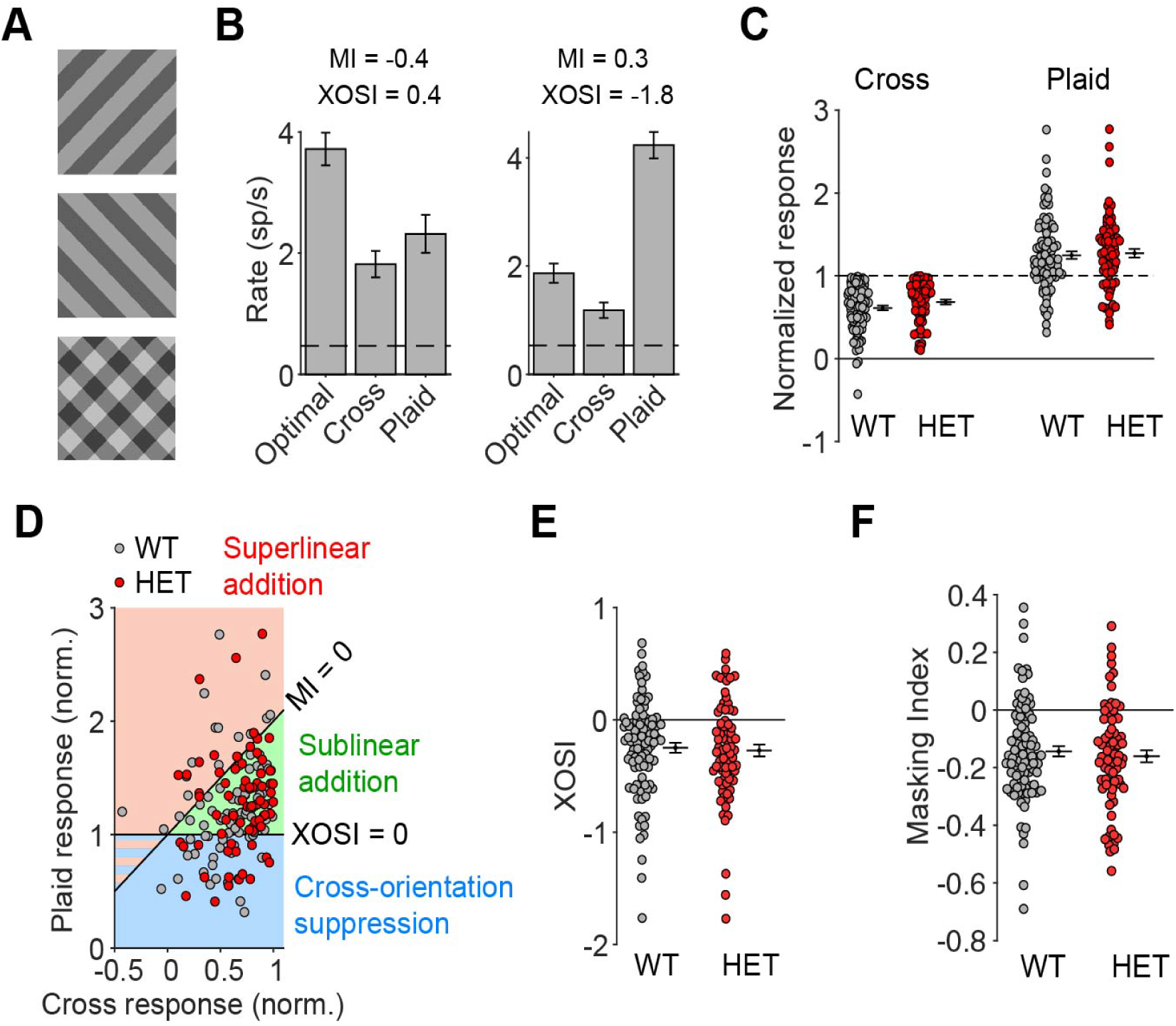
No difference in cross-orientation suppression in Vglut2-Stxbp1 HET vs. WT mice. **(A)** Example stimuli. Two single orientations at 50% contrast and one combination plaid. **(B)** Example of responses of a neuron showing cross-orientation suppression (left) and of a neuron showing superlinear addition (right). Bars indicate mean ± s.e.m. **(C)** Population responses to the cross-oriented and plaid stimulus, normalized to the response to the optimal grating. Responses do not differ between Vglut2-Stxbp1 WT and HET mice. Dots indicate neurons (7 WT mice, 5 HET mice). Error bars indicate mean ± s.e.m. **(D)** Responses, normalized to the response to the optimal grating, for plaid versus cross-oriented gratings. The lines for linear addition (MI = 0) and no cross-orientation suppression (XOSI = 0) are shown. Zones indicating superlinear addition, sublinear addition and cross-orientation suppression are displayed in red, green, and blue, respectively. **(E)** No difference in distribution of XOSI between Vglut2-Stxbp1 WT and HET mice. **(F)** No difference in the masking index. Detailed statistics are in **Supp. Table 1**.

So far, we have shown that Vglut2-Stxbp1 HET mice have a shorter V1 response half-life than their WT siblings, but that the receptive fields, contrast normalization and cross-orientation suppression are similar between the two genotypes. There was, however, a notable difference in the amount of direction selectivity between the Vglut2-Stxbp1 WT and HET mice (**Fig. 6A-B**). This was expressed both by a significantly lower direction selectivity index (DSI) in V1 neurons of HET mice (**Fig. 6C,** p = 0.003), as well as in a lower number of directive selective neurons (with DSI>0.33, **Fig. 6D,** p = 0.004). These results are consistent with work that shows short-term synaptic depression could cause or have an influence on direction selectivity in the visual cortex (Buchs & Senn, 2002; Chance et al., 1998). It is also consistent with results that direction selectivity in mouse primary visual cortex can emerge through the convergence of transient-firing and sustained-firing thalamocortical afferents (Lien & Scanziani, 2018). These findings are consistent with the idea that shortening the response half-life of thalamocortical input could impede the cortical summation that is necessary for direction selectivity to emerge, although effects at other Vglut2-expressing synapses or developmental changes could also contribute.

**Figure 6.**
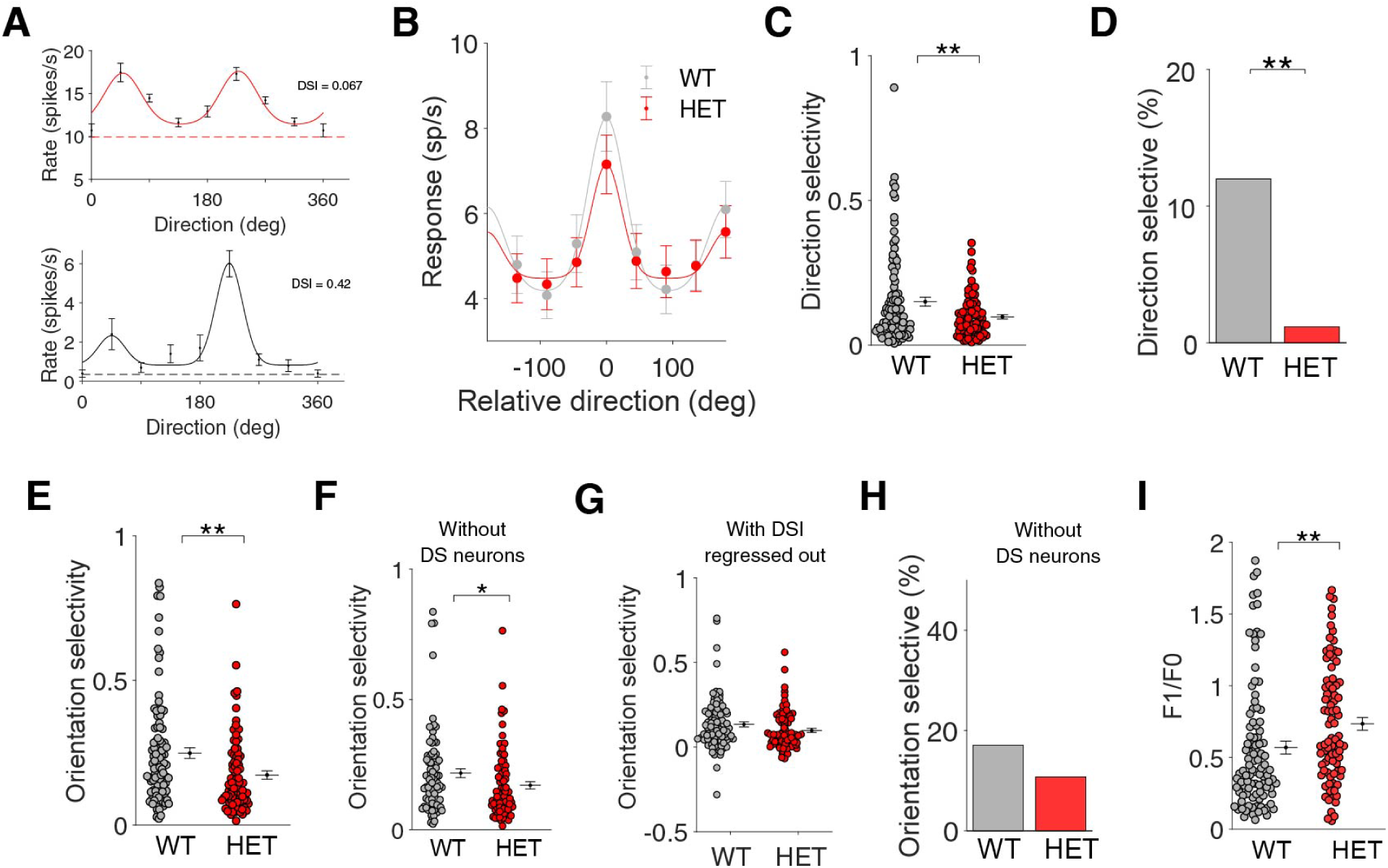
Direction selectivity in V1 is reduced in Vglut2-Stxbp1 HET mice. **(A)** Example direction tuning of neuron in Vglut2-Stxbp1 HET (top) and WT mouse (bottom). Error bars indicate mean ± s.e.m. **(B)** Population mean response for each direction relative to the preferred direction (WT: 100 units from 7 mice, HET: 85 units from 5 mice). **(C)** Direction selectivity is significantly reduced in Vglut2-Stxbp1 HET mice. Error bars indicate mean ± s.e.m. Dots indicate units. **: p<0.01. **(D)** Number of direction-selective (DSI>0.33) units is significantly lower in Vglut2-Stxbp1 HET mice. **: p<0.01. **(E)** Orientation selectivity is also significantly reduced in Vglut2-Stxbp1 HET mice. **: p<0.01. **(F)** Orientation selectivity is still significantly reduced if only non-direction-selective units are considered. *: p<0.05. **(G)** When the correlation with the direction selectivity is regressed out, there is a trend for lower orientation selectivity in Vglut2-Stxbp1 HET mice. **(H)** Number of specifically orientation selective cells (OSI>0.33) is not significantly reduced. **(I)** F1/F0 phase sensitivity is significantly higher in Vglut2-Stxbp1 HET mice. **: p<0.0. Detailed statistics are in **Supp. Table 1**.

The V1 neurons of the HET mice also showed a significantly lower orientation selectivity index (OSI, **Fig. 6E,** p = 0.001). However, orientation and direction selectivity are intrinsically linked: cells that are strongly direction-selective will also be orientation selective (but not the other way around). Therefore, we attempted to tease apart the effect on the DSI and the OSI by removing the DSI effects from the OSI results. When removing all direction-selective neurons (DSI>0.33) from the OSI analysis, the difference between the WT and HET mice became smaller (**Fig. 6F**, p = 0.04) and, after regressing out the correlation of the direction selectivity from the orientation selectivity index, the difference was no longer significant (**Fig. 6G,** p = 0.065). Similarly, the number of orientation selective neurons was not significantly different between WT and HET mice when removing the direction-selective neurons (**Fig. 6H,** p = 0.23).

Finally, the V1 responses of the Vglut2-Stxbp1 HET mice to the drifting gratings were notably more modulated at the drift frequency than those of the WT mice. Cells that are modulated at the drift frequency are usually called ‘simple’ cells, whereas cells that are less modulated are called ‘complex’ cells - referring to the fact that their responses do not linearly follow from their receptive field structure (Hubel & Wiesel, 1962; Skottun et al., 1991). Whereas we recorded mostly complex cells from the Vglut2-Stxbp1 WT mice, V1 of the HET mice appeared to be dominated more by simple cells. The difference in the F1/F0, a measure for response linearity, was significant (**Fig. 6I**, p = 0.0097). As complex cell responses are assumed to originate from the integration of input from cells with a more simple response (Hubel & Wiesel, 1962), this indicates that neural integration is affected in the Vglut2-Stxbp1 HET mice.

## Discussion

In this study, we used a Vglut2-Cre-dependent heterozygous deletion of Stxbp1 to examine how increased short-term depression affects visual processing in mouse V1. We confirmed increased short-term depression in white-matter inputs to layer 4 and found that visual responses were shorter in Vglut2-Stxbp1 HET mice than in WT mice. Despite this clear alteration in response dynamics, contrast saturation and cross-orientation suppression were unchanged. By contrast, direction selectivity was reduced and responses to drifting gratings were more strongly modulated at the stimulus frequency. These findings dissociate divisive normalization from other aspects of visual response integration: altered short-term plasticity changes the temporal dynamics and feature selectivity of V1 responses, but does not measurably affect the two forms of divisive normalization tested here.

In the heterozygous Cre-dependent knock-out mice, we did find differences in the direction and orientation selectivity, as well as in the linearity (F1/F0) of the V1 responses. However, we used a Vglut2-Cre line to knock-out Stxbp1 heterozygously in thalamic relay neurons (Vong et al., 2011) as Vglut2 is the primary vesicular glutamate transporter in the thalamus. Although Vglut2 is only sparsely present in adult V1 (Lein et al., 2007), there is expression of Vglut2, and therefore of Cre-recombinase, during development in both the retina and the visual cortex (Oh et al., 2014). We can therefore not rule out that visual processing at other stages than the thalamocortical synapse is also affected, and that these changes in direction and orientation selectivity are caused by changes in other synapses than the thalamocortical synapses. Regardless of this, we did not find an effect of the knockout on contrast tuning and cross-orientation suppression, suggesting that any unforeseen effects of the knockout also did not affect the mechanisms behind these phenomena and therefore did not affect our conclusion regarding divisive normalization.

Our experiments provide new evidence in the search for the mechanism(s) behind divisive normalization. More specifically, our results reject the idea that short-term depression is necessary for gain control and divisive normalization in response to visual stimuli (Abbott et al., 1997; Carandini et al., 2002). Given that STD is so commonly found in the thalamocortical synapse (Bannister et al., 2002; Kloc & Maffei, 2014; Stratford et al., 1996) and the clear modeling predictions, it is surprising that an increased STD at this synapse has so little influence on the effects of normalization. Previous work suggested that ongoing activity can tonically depress thalamocortical synapses in vivo, thereby limiting the additional depression evoked by sensory stimulation (Boudreau & Ferster, 2005; Reig et al., 2006). However, these recordings were in anesthetized animals while recordings in the awake rabbit showed that stimulation can still result in strong STD at the thalamocortical synapse (Swadlow et al., 2002). The influence of thalamocortical depression also depends on brain state: in awake mice, V1 responses can remain tightly locked to thalamic input, whereas under anesthesia thalamocortical synaptic depression becomes rate-limiting and disrupts the temporal fidelity of sensory transmission (Reinhold et al., 2015). One way to reconcile these findings with STD-based models is that the effect of short-term depression on response gain may saturate at physiologically relevant firing rates, such that further increases in depression have only limited impact on steady-state responses (Kayser et al., 2001).

Divisive normalization is a computational description of response scaling rather than a single circuit mechanism, and it has been used to describe responses across sensory brain areas and behavioral contexts (Bonin et al., 2005; Carandini & Heeger, 2012; Heeger, 1992; Ohshiro et al., 2017; Solomon & Kohn, 2014). Accordingly, short-term depression is only one of several proposed biological mechanisms that could implement normalization in V1. Another proposed mechanism in V1 is GABAergic inhibition (Carandini et al., 1997; Ferguson & Cardin, 2020; Keller & Martin, 2015), although experiments testing this hypothesis in cat V1 using GABA_A_-blockers saw a change in response gain only, and not in contrast tuning or cross-orientation suppression (Katzner et al., 2011). In mouse V1, distal cortical activation can produce divisive normalization through a reduction in recurrent excitation rather than an increase in inhibition, suggesting that cortical normalization may depend on the dynamics of recurrent excitatory-inhibitory networks (Sato et al., 2016). This interpretation is also consistent with models in which normalization-like suppression arises in inhibition-stabilized cortical networks with strong recurrent excitation (Rubin et al., 2015). For cross-orientation suppression specifically, however, recent work suggests a possible thalamic origin (Barbera et al., 2022). The latter study was very suggestive that the origin of XOS in V1 was lying in the selectivity of the thalamic input, but it could not rule out a role for STD in the thalamocortical synapse.

Our results did not support the proposed role of short-term depression in divisive normalization. Instead, they point to a role in shaping the temporal persistence and integration of visual responses. Short-term plasticity is often described as a temporal filter: depressing synapses preferentially transmit response onset and low-frequency changes while attenuating sustained or rapidly repeated input (Abbott & Regehr, 2004; Citri & Malenka, 2008; Fortune & Rose, 2001). Consistent with this interpretation, increased short-term depression in the Vglut2-Stxbp1 HET mice shortened visually evoked responses without reducing their peak amplitude. Such changes in response duration could alter the temporal overlap between converging inputs and thereby affect the construction of feature selectivity, even when steady-state contrast normalization remains intact.

Our results regarding the reduced direction selectivity in the Vglut2-Stxbp1 HET mice corroborate research suggesting that de novo generation of direction selectivity in V1 comes about by the summation of transient and sustained thalamocortical responses (Lien & Scanziani, 2018; Saul & Humphrey, 1990; Stanley et al., 2012): if a drifting grating activates first a neuron with a sustained response, and then a neuron with a transient response, their summed outputs would be higher than if they were activated in the opposite order. Our findings suggest that our knockout model decreases the response half-life at the thalamic relay cell synapses, thereby impeding the cortical summation that is necessary for direction selectivity to emerge. However, the decrease in orientation selectivity cannot be explained by the transient/sustained model. As V1 orientation selectivity in principle can be explained by the summation of inputs from neurons with neighboring receptive fields (Hubel & Wiesel, 1962), no temporal integration should be needed for orientation selectivity to emerge. But even when removing the direction selectivity effect from the orientation selectivity effect (**Fig. 6**), a reduction in orientation selectivity appears to persist. This result, as well as the increased linearity (F1/F0) of the cortical responses, raises questions about precisely how the Vglut2-Stxbp1 knockout model alters (thalamo-)cortical integration. More experiments would be needed to determine whether these changes involve altered thalamocortical response timing, intracortical integration, or circuit mechanisms such as the spatial organization of excitatory and inhibitory inputs that has been linked to direction selectivity in mouse V1 (Rossi et al., 2020).

In conclusion, increasing short-term depression through heterozygous Stxbp1 deletion shortened visual responses and altered direction selectivity and response linearity, while leaving contrast saturation and cross-orientation suppression unchanged. These results argue against short-term depression at Vglut2-expressing inputs as the principal mechanism underlying these forms of divisive normalization in mouse V1. At the same time, they support a role for short-term plasticity in determining the temporal window over which visual inputs are integrated and transformed into selective cortical responses. Short-term depression may therefore shape when and how visual signals are combined, rather than providing the normalization operation itself.

## Materials & Methods

### Experimental animals

All mice came from the crossing of a homozygous Vglut2-ires-Cre parent (Slc17a6tm2(cre)Lowl/J, Jax, #016963; (Vong et al., 2011)) and a heterozygous LoxP-Stxbp1 parent (Heeroma et al., 2004). For the slice experiments, we used a total of 10 mice (4 HET, 6 WT; 6 male, 4 female). Mice were 21-42 days old at the experiment date. For the in vivo experiments, we used a total of 14 mice (7 HET, 7 WT; 9 male, 5 female). Mice were 60-100 days old at the experiment date. For each Vglut2-Stxbp1 HET mouse, we used a Vglut2-Stxbp1 WT sibling as control. The mice were housed in pairs or groups in a regular light/dark cycle (12/12 hr) with ad libitum access to food pellets and water. All experimental protocols were approved by the institutional animal care and use committee of the Royal Netherlands Academy of Sciences (KNAW) and were in accordance with the Dutch Law on Animal Experimentation.

### Slice electrophysiology

Mice were anesthetized with 5% isoflurane, directly followed by decapitation and the brain was quickly removed from the skull and placed into ice-cold slicing artificial CSF (aCSF) of the following composition (in mM): 125 NaCl, 3 KCl, 25 glucose, 25 NaHCO3, 1.25 Na2H2PO4, 1 CaCl2, 6 MgCl2, 1 kynurenic acid, saturated with 95% O2 and 5% CO2, pH 7.4). Coronal slices (350 μm) containing the primary visual cortex were cut with a Vibratome (1200S, Leica Microsystems) and brain slices were allowed a recovery period at 35°C for 35 min, thereafter stored at room temperature. Slices were transferred to an upright microscope (BX51WI, Olympus Nederland) equipped with oblique illumination optics (WI-OBCD; numerical aperture, 0.8). Neurons were visualized using either 4× (0.80W) or 60× (1.00W) water-immersion objectives (Olympus). The microscope bath was perfused with oxygenated (95% O2, 5% CO2) aCSF consisting of the following (in mM): 125 NaCl, 3 KCl, 25 D-glucose, 25 NaHCO3, 1.25 Na2H2PO4, 2 CaCl2, and 1 MgCl2. Whole cell patch clamp recordings were made in layer 4 neurons with a bipolar stimulating electrode in white matter below the patched neurons. Layer 4 neurons were identified by the density of cell neocortical layer boundaries in contrast to other layers and the small soma that characterize this layer, especially in contrast to layer 5 pyramidal neurons with their typical large triangular shape (Scala et al., 2019). The layer identity of each neuron was also confirmed by the visualization of their position after the post hoc biocytin staining. Whole cell voltage-clamp recordings were made using Axopatch 200B (Molecular Devices) and patch pipettes were pulled from borosilicate glass (Harvard Apparatus) pulled to an open tip resistance of 2–5 MΩ (whole cell). Patch pipettes were filled with (in mM) the following: 130 K-gluconate, 10 KCl, 4 Mg-ATP, 0.3 Na2-GTP, 10 HEPES, and 10 Na2-phosphocreatine, pH 7.4, adjusted with KOH, 280 mOsmol/kg, to which 5 mg/ml biocytin was added. Currents were evoked from the bipolar stimulating electrode using an ISO-Flex stimulus isolator (A.M.P.I.) in the white matter of the cortex to stimulate the thalamocortical tracts. Patched cell were given ten sets of ten pulses, administered with three different frequencies for each cell: 10 Hz, 25 Hz and 40 Hz. For each frequency, the amplitude of the stimulation pulses was increased in for each set with 20 mA increments, ranging from 40 mA to 100 mA. These different frequencies were applied consecutively, with approximately 30 seconds of rest between each set. All recordings were made at 34°C. Voltage was analog low-pass filtered at 5 kHz (Bessel) and digitally sampled at 50 kHz – 100 kHz using an analog to digital converter (ITC-18, HEKA Electronic) and data acquisition software AxoGraph X (v.1.7.6 and v.1.8). Both fast and slow capacitances were fully compensated. Series resistance throughout the recordings was monitored under voltage clamp by applying a hyperpolarizing square pulse (−10 mV for 10 s) and compensated. The series resistance was typically < 21 MΩ. During recording, a holding potential of −70 mV was used. All potentials were corrected for liquid junction potential (LJP = 14.1 mV), using Clampex. After patching a neuron, we applied a step current to observe the spiking response to differentiate between inhibitory and excitatory neurons, following the electrophysical description of layer 4 of mouse visual cortex (Kloc & Maffei, 2014; Scala et al., 2019). The excitatory neurons show large action potentials width, high action potential amplitude and shallow afterhyperpolarization (Scala et al., 2019).

### Analysis of slice electrophysiology

Individual traces (between 500 and 1500ms duration) were filtered with a Gaussian blur filter of 1 KHz. The EPSC recordings were analyzed using event detection from Axograph and a Matlab script with representatives EPSC template, unique for the recording file. Detected events were averaged for further analysis of peak amplitude. The responses were selected when the median EPSC amplitude over all pulses for one cell must be larger than 2x the root mean square (RMS). The EPSCs were considered monosynaptic if their delay from stimulus onset was <2.5 ms and did not show any additional peak on the decay phase of the postsynaptic current (Kloc & Maffei, 2014). All neurons showing disynaptic responses, or with a resting membrane potential below −80 mV or above -55 mV were excluded from the analysis.

### Immunohistochemistry for slice electrophysiology

Layer 4 neurons were filled with 5 mg/mL biocytin during whole-cell patch clamp recordings for at least 20 min. Slices were fixed for 20 min with 4% paraformaldehyde (PFA). The slices were then kept in PBS at 4°C. Acute slices were blocked with 5% normal goat serum (NGS) and 2% Triton X-100 for 2h at room temperature on shaker plate. For biocytin-labelled cells, streptavidin biotin-binding protein (Streptavidin Alexa 633, 1:500, Invitrogen, RRID:AB_2315383) was diluted in 5% NGS and 2% Triton X-100 and incubated overnight at room temperature on shaker plate. Finally, sections were mounted on glass slides and cover slipped with Vectashield H1000 fluorescent mounting medium and sealed. Images were collected with a Leica SP8 X (DM6000 CFS; acquisition software, Leica Application Suite AF v3.2.1.9702) confocal laser-scanning microscopes (Leica Microsystems). Confocal images were acquired using 40× (numerical aperture, 1.3) oil-immersion objectives. Only neurons located in layer 4 were used in the analysis.

### In vivo electrophysiology

Mice were anesthetized by an intraperitoneal (IP) injection of 1.2 g urethane per kg body weight, supplemented by an IP injection of 8 mg chlorprothixene per kg body weight. We injected atropine sulfate (0.1 mg per kg, s.c.) to reduce mucous secretions. Additional doses of urethane were injected when a response to a toe-pinch was observed. Temperature was measured with a temperature probe underneath the mouse and maintained by a feedback-controlled heating pad set to 36.5 degrees. We cleared the throat so the mouse could breathe freely and then head fixed the mouse by ear and bite bars. We trimmed the hair on the head and applied lidocaine spray on the skin for local analgesia. We then made an incision in the skin and drilled a ∼1 mm craniotomy for the electrode over V1 at 0.6 mm anterior and 2.9 mm lateral from the lambda cranial landmark. For the reference site, we drilled a 0.5 mm craniotomy in the contralateral frontal bone plate. We first inserted the reference wire into the reference site. We recorded using a laminar silicon electrode (A1 × 16-5mm-50-177-A16, 16 channels spaced 50 μm apart, Neuronexus). For some mice, the electrode was coated with a green neuro-DiO (Biotium) or a red diI (Fisher Scientific) tracer solution for later histological verification of the recording site. Then the electrode was inserted into the brain to a depth of ca. 800 µm. We used black tin foil (Thorlabs) to reduce external noise on the electrode. We waited at least 15 minutes for the brain to stabilize before starting the recording. The electrical signal from the electrodes was amplified and sampled at 24.4 kHz using a Tucker-Davis Technologies recording system. If, after recording all the visual stimuli, the anesthesia of the mouse was still stable, we removed the electrode from the brain, and repeated the experiment at a slightly different recording location, typically 100-200 μm away from the previous site.

### Visual stimulation

Visual stimuli were projected onto a back-projection screen placed 15 cm from the mouse with a PLUS U2-X1130 DLP projector (gamma-corrected, mean luminance = 40.6 cd/m2). The size of the projection was 76 cm by 56 cm, the field-of-view was 136° × 101.6°, the resolution was 1024 × 768 pixels, and the refresh rate was 60 Hz. In order to assess the receptive field location of the recorded sites, we presented a 5 min movie (5 frames per second) of small white squares, approximately 10 degrees wide, in random positions on a white background (ratio of white to black area: 1:34) (Heimel et al., 2010).

For all further stimuli, we used full-screen square wave gratings with a spatial frequency of 0.02 cycles per degree and a temporal frequency of 2 Hz. First, we mapped the direction tuning of the recorded sites using 100% contrast gratings. We mapped 8 different directions and displayed each stimulus 8 times. Both the stimulus duration and the intertrial interval (ITI) were 3 seconds. We next mapped the contrast response curve using the preferred grating direction of the recorded site, defined as the direction that evoked the largest multi-unit response during the direction-tuning stimulus. We displayed grating contrasts of 30, 50, 70 and 95%, and repeated each stimulus 20 times. Stimulus duration and ITI were 3 s. Finally, we recorded cross-orientation suppression. For this, we combined 50% contrast drifting gratings at the site-preferred direction and the orthogonal direction. The combination was a linear sum of the difference of each of the gratings compared to the mean luminance, added to the mean luminance. The result is a 100% contrast plaid. Combining the absence/presence of each grating yielded 4 stimuli: grey screen, optimal grating, orthogonal grating and plaid. Again, stimulus duration and ITI were 3 s and each stimulus was repeated 20 times.

### Histology for in vivo experiments

After the recording, the mice were deeply anesthetized with Nembutal and transcardially perfused with phosphate-buffered saline (PBS), followed by 4% paraformaldehyde (PFA) in PBS. We extracted the brain and post-fixated it overnight in 4% PFA before moving it to a PBS solution. We cut the brains into 75-um-thick coronal slices and mounted them on glass slides in Vectashield DAPI solution (Vector Laboratories). We imaged the slices on a Zeiss Axioscan.Z1 using ZEN software. The resulting histology images were used to confirm the location of the electrode.

### Analysis of in vivo data

All offline analysis was performed using MATLAB (R2019a; MathWorks). The analysis scripts are available at https://github.com/heimel/Stxbp1_KO. For analysis of the extracellular recordings, we use the InVivoTools toolbox, available online at https://github.com/heimel/InVivoTools. We incorporated Kilosort 2.0 in this code (https://github.com/MouseLand/Kilosort/releases/tag/v2.0; (Stringer et al., 2019)) to preprocess the raw data, using a 2x standard deviation threshold to isolate spikes. Spike were sorted automatically using KlustaKwik (https://klusta-team.github.com/klustakwik; (Kadir et al., 2014) and only clusters were selected for which the spike height was at least 5 times larger than the standard deviation of the deviation of the assigned spikes to their mean cluster template . The spike rate was computed by binning the spikes over all trials in 10 ms bins. For presentation purposes, the spike rate in the histograms was smoothed by convolution with a Gaussian with a 20 ms width. The response was calculated by subtracting the spontaneous rate, which was the mean rate in the period starting 0.5 s after the stimulus offset and ending at stimulus onset. For the analysis of the receptive field stimulus, we computed the spike-triggered average. We only included cells that were responsive, which we determined by setting a threshold of 0.95 for the area under the curve of an ROC analysis for detection of the preferred white patch in the spike-triggered average. All patches that had a mean spike-triggered average more than 3 times the standard deviation above the mean luminance of the stimulus were considered part of the on-receptive field. The peak response was measured during presentation of the preferred white patch in the spike-triggered average. The response half-life was the amount of time after the peak response for the response to fall below half the peak response. For the tests with gratings or plaids, we used as inclusion criteria that units needed to be responsive, as measured by a p-value on the Zeta test below 0.05 (Montijn et al., 2021) and have a minimum response of 2 spikes/s during the stimulus presentation for at least one of the different stimuli. For computing the C50 and dynamic range, we fitted the responses for the different contrasts for each unit with a Naka-Rushton curve (Albrecht & Hamilton, 1982) and computed the values from the fits. The C50 was the contrast at half the maximum response. The dynamic range was the range of contrast between a quarter and three-quarters of the maximum response. For the plaid stimuli, we computed the cross-orientation suppression index (XOSI) as XOSI = 1 – Response to plaid / Response to optimal grating. The masking index (MI) was computed as the (Response to plaid - Response to optimal grating - Response to orthogonal grating) / (Response to plaid + Response to optimal grating + Response to orthogonal grating) (Barbera et al., 2022; Guan et al., 2020). The OSI was defined as OSI=√(∑R(φ) sin(2φ)² + ∑R(φ) cos(2φ)²) / ∑R(φ), where φ is the direction of the stimulus and R(φ) the neuron’s response. This is equal to 1–circular variance. DSI was defined by DSI=√(∑R(φ) sin(φ)² +∑R(φ) cos(φ)²) /∑R(φ).

### Experimental design and statistical analysis

Based on the published slice recordings with heterozygous knock-out of Stxbp1 and previous success rate of in vivo recordings, we performed a power analysis before the experiments to determine the number of animals. For all comparisons of numerical data between groups of neurons from HET and WT mice, we used two-tailed, two-sample Student t-tests if the data was normal and the Mann-Whitney U test if the data failed a Shapiro-Wilk normality test. For comparison of the EPSC size during the pulse trains (**Fig. 2G**), we use a non-linear mixed effects model, with the model a + (1-a) exp( - (pulse #-1)/b) where a has a fixed effect per genotype (HET/WT) and a random effect per cell. A Wald z-test was used to test the effect of genotype. For categorical data, chi^2^ tests were used.

## Acknowledgements

We are thankful to Christiaan Levelt for connecting our research interests.

