## Supplementary information for "Divisive normalization in mouse V1 is unchanged by increased thalamocortical short-term depression"

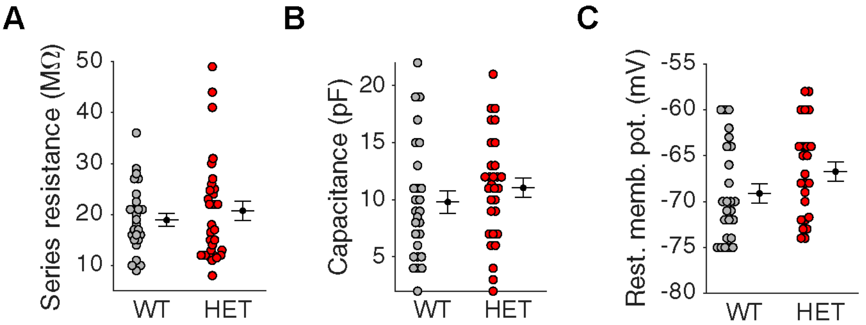


**Supplementary Figure 1. No difference in cell intrinsic properties in Vglut2-Stxbp1 HET vs. WT mice.** **(A)** No difference in series resistance. **(B)** No difference in capacitance. **(C)** No difference in resting membrane potential.

### Supplementary Table 1 with statistics

Figure 2

| Panel | Comparison | Mean & s.e.m. | Units | Test | Test Statistic | p-value |
|---|---|---|---|---|---|---|
| 2E | WT vs HET | WT: 0.99 ± 0.15 nA<br>HET: 1.11 ± 0.17 nA | 23<br>23 | Mann-Whitney | U = 512 | 0.54 |
| 2F | 10 Hz offset: WT vs HET | WT: 0.88 ± 0.04<br>ΔHET: -0.12 ± 0.06 | 23<br>23 | Non-linear mixed effects model, Wald test | z = -2.09 | 0.037 |
| | 25 Hz: WT vs HET | WT: 0.88 ± 0.04<br>ΔHET: -0.20 ± 0.06 | 23<br>23 | Non-linear mixed effects model, Wald test | z = -3.21 | 0.0013 |
| | 40 Hz: WT vs HET | WT: 0.84 ± 0.06<br>ΔHET: -0.21 ± 0.08 | 23<br>23 | Non-linear mixed effects model, Wald test | z = -2.60 | 0.0094 |
| 2G | 10 Hz: WT vs HET | WT: 0.15 ± 0.04<br>HET: 0.26 ± 0.04 | 24<br>24 | Mann-Whitney | U = 493 | 0.051 |
| | 25 Hz: WT vs HET | WT: 0.12 ± 0.05<br>HET: 0.37 ± 0.04 | 23<br>23 | Mann-Whitney | U = 369 | 0.00017 |
| | 40 Hz: WT vs HET | WT: 0.19 ± 0.08<br>HET: 0.40 ± 0.08 | 23<br>23 | Mann-Whitney | U = 431 | 0.017 |

Figure 3

| Panel | Comparison | Mean & s.e.m. | Units | Test | Test Statistic | p-value |
|---|---|---|---|---|---|---|
| 3C | WT vs HET | WT: 34.9 ± 1.3 deg<br>HET: 34.8 ± 1.3 deg | 53<br>44 | Two-sample t-test | t[95] = 0.06 | 0.95 |
| 3F | WT vs HET | WT: 9.3 ± 1.1 Hz | 53 | Two-sample | t[95] = -1.24 | 0.22 |

| | | | | | | |
|---|---|---|---|---|---|---|
| | | HET: 10.8 ± 1.1 Hz | 44 | t-test on log transform | | |
| 3G | WT vs HET | WT: 0.118 ± 0.008 s<br>HET: 0.089 ± 0.006 s | 53<br>44 | Two-sample t-test | t[95] = 2.90 | 0.0047 ** |

Figure 4

| Panel | Comparison | Mean & s.e.m. | Units | Test | Test Statistic | p-value |
|---|---|---|---|---|---|---|
| 4D | WT vs HET | WT: 40.6 ± 1.7 %<br>HET: 36.8 ± 1.7 % | 65<br>55 | Two-sample t-test | t[118] = 1.59 | 0.11 |
| 4E | WT vs HET | WT: 33.9 ± 0.9 %<br>HET: 34.2 ± 1.2 % | 65<br>55 | Two-sample t-test | t[118] = -0.22 | 0.83 |

Figure 5

| Panel | Comparison | Mean & s.e.m. | Units | Test | Test Statistic | p-value |
|---|---|---|---|---|---|---|
| 5C | Cross: WT vs HET | WT: 0.61 ± 0.03<br>HET: 0.68 ± 0.02 | 93<br>73 | Two-sample t-test | t[164] = -1.75 | 0.082 |
| | Plaid: WT vs HET | WT: 1.25 ± 0.04<br>HET: 1.27 ± 0.05 | 93<br>73 | Two-sample t-test | t[164] = -0.34 | 0.74 |
| 5E | WT vs HET | WT: -0.25 ± 0.04 %<br>HET: -0.27 ± 0.05 % | 93<br>73 | Two-sample t-test | t[164] = 0.34 | 0.73 |
| 5F | WT vs HET | WT: -0.14 ± 0.02 %<br>HET: -0.16 ± 0.02 % | 93<br>73 | Two-sample t-test | t[164] = 0.62 | 0.54 |

Figure 6

| Panel | Comparison | Mean & s.e.m. | Units | Test | Test Statistic | p-value |
|---|---|---|---|---|---|---|
| 6C | WT vs HET | WT: 0.150 ± 0.015<br>HET: 0.098 ± 0.008 | 100<br>85 | Two-sample unequal variances t-test | t[144.8] = 3.0 | 0.0031 ** |
| 6D | WT vs HET | WT: 12 %<br>HET: 1 % | 100<br>85 | $Chi^2$ test | - | 0.0041 ** |
| 6E | WT vs HET | WT: 0.249 ± 0.018<br>HET: 0.173 ± 0.014 | 100<br>85 | Two-sample unequal var. t-test | t[178.1] = 3.3 | 0.0013 ** |
| 6F | WT vs HET | WT: 0.218 ± 0.017<br>HET: 0.171 ± 0.014 | 88<br>84 | Two-sample t-test | t[170] = 2.1 | 0.037 * |
| 6G | WT vs HET | WT: 0.134 ± 0.015<br>HET: 0.099 ± 0.012 | 100<br>85 | Two-sample unequal var. t-test | t[180.1] = 1.9 | 0.065 |
| 6H | WT vs HET | WT: 17 %<br>HET: 11 % | 88<br>84 | $Chi^2$ test | - | 0.23 |
| 6I | WT vs HET | WT: 0.569 ± 0.045<br>HET: 0.735 ± 0.044 | 100<br>85 | Two-sample t-test | t[183] = -2.6 | 0.0097 ** |
